# The Performance and Potential of Deep Learning for Predicting Species Distributions

**DOI:** 10.1101/2024.08.09.607358

**Authors:** Benjamin Kellenberger, Kevin Winner, Walter Jetz

## Abstract

Species distribution models (SDMs) address the whereabouts of species and are central to ecology. Deep learning (DL) is poised to further elevate the already significant role of SDMs in ecology and conservation, but the potential and limitations of this transformation are still largely unassessed.

We evaluate DL SDMs for 2,299 terrestrial vertebrate and invertebrate species at continental scale and 1km resolution in a like-for-like comparison with latest implementation of classic SDMs. We compare two DL methods (a multi-layer perceptron (MLP) on point covariates and a convolutional neural network (CNN) on geospatial patches) against existing SDMs (Maxent and Random Forest). On average, DL models match, but do not surpass, the performance of existing methods. DL performance is substantially weaker for species with narrow geographic ranges, fewer data points, and those assessed as threatened and hence often of greatest conservation concern. Furthermore, information leakage across dataset splits substantially inflates performance metrics, especially of CNNs. We find current DL SDMs to not provide significant gains, instead requiring careful experimental design to avoid biases. However, future advances in DL-supported use of ancillary ecological information have the potential to make DL a viable instrument in the larger SDM toolbox. Realising this opportunity will require a close collaboration between ecology and machine learning disciplines.

## 1 Introduction

Ever-changing conditions of our biosphere make the geographic understanding of biodiversity at a large scale ever more pressing [6]. For predicting patterns and capturing potential drivers of spatial biodiversity dynamics species distribution models (SDMs) have become the central framework and toolbox [84]. In SDMs, the geospatial locations on Earth where species have been observed are related to characteristics of their immediate environment, such as climatic conditions or soil characteristics [22]. SDMs have become a staple approach for ecological modelling and have been used to approximate the distribution of lesser-known taxa [41], map the spread of pathogens through the distribution of hosts [92], assess human-wildlife coexistence potential [85], model the risks of invasive species [79], and forecast species’ adaptation potential to changing environmental conditions [16]. Over the past decade, SDMs have seen immense use in tens of thousands of publications based on a range of methodologies (*e.g.*, [30, 64, 58, 23, 66, 70]). While most are single-species models, some studies have also included joint SDMs which model the distribution of multiple species together [69], with potential, albeit debated [27], ability to capture additional ecological mechanisms.

Recently, a new generation of SDMs has emerged based on deep learning [46] (DL), able to model arbitrarily complex relationships [34], for larger dataset corpora [17], and a very high number of species at once [18, 51]. These models have gained rapid momentum with high reported performances [10, 17, 26], which might suggest they could in the future overtake traditional, “shallow” SDMs.

### 1.1 Limitations of existing SDMs

DL might offer the opportunity to overcome some of the hard limitations seen in current SDMs and catalyse a next generation of biodiversity analytics [10, 11, 14]. Given the complex nature of species-environment relationships, existing, conventional methodologies are often necessarily limited in their expressivity and scalability. Key limitations of current SDMs include: *(i.)* unrealistic assumptions, *e.g.* that relatively simple, *a priori* model specifications are able to address complex relationships or interactions with their abiotic and biotic setting [87]; *(ii.)* ignorance of spatial context (*i.e.*, environmental characteristics are commonly only sampled for the immediate centre of observations, discarding potentially vital signals from the landscape surroundings [47, 48]; *(iii.)* inability to leverage ancillary information about species ecological attributes or relationships beyond environmental descriptors; and *(iv.)* increasing inability to process and use the growing amounts and types of ecological data emerging today.

A growing number of sophisticated sensors on earth orbiting satellites or aerial vehicles deliver environmental covariates at increasingly fine resolution and large extents [45], and species occurrence records have accumulated dramatically in absolute quantities as well as number of species covered [51], albeit sometimes without growth in actual coverage [60]. While joint SDMs can process such growing multi-species data in a single framework [69, 63], they typically require a completely defined observation space, with presence and absence of all species in question known for every sampled location. Such scheme can only be obtained by expensive (and hence small-scale) systematic surveys, or aggregation of observations in space at the cost of resolution. Joint SDMs usually also involve computationally expensive training schemes limiting their scalability. Neither property makes them amenable to present-day citizen science-derived occurrence records, which are large in quantities and form a highly incomplete observation space where the probability of presence is only known for a very low number of species for each geographic location. To date, effective avenues for modelling the geographic distribution of many species jointly is missing.

### 1.2 Deep Learning (DL) applied to Species Distributions

The last decade has seen significant development in the field of deep learning [46] (DL), which has the potential to address the above shortcomings of existing SDMs. DL represents a family of modular machine learning methodologies that learn both decision functions and features for the prediction task(s) at hand in a layer-wise fashion [28]. DL models have significantly reduced the prediction error in a vast number of applications, including image classification [44, 33], natural language processing [62], remote sensing [93], as well as ecology-related tasks like animal detection in camera trap [59] and aerial images [42, 83]. In wake of the big data revolution that has also reached ecology, DL offers promising prospects for applications like SDMs.

As seen in other areas related to machine learning, an initial body of work used feed-forward (artificial) neural networks for species distribution modelling (*e.g.*, [53, 49, 31, 90, 9, 89]). *Sensu stricto*, these feed-forward neural networks do belong to the family of DL models as their fundamental working principle (prediction by sequential application of layers) and training procedure (gradient descent by backpropagation [73]) are similar to modern-day equivalents. However, they only use a small number (*e.g.*, two to three) of linear, resp. fully-connected layers, thus have relatively few learnable parameters, and have generally been used as a drop-in replacement for shallow machine learning models. As such, they do not scale particularly well to large datasets and do not include recent DL discoveries like skip connections [33], batch normalisation [36], or dropout [78]. Also, none of them are convolutional or include spatial context in any other form.

Recent research has introduced many concepts and changes that set modern-day deep learning models apart from these earlier models. One of the first studies to also investigate CNNs for SDM is Botella et al. [10], who pitch a (still relatively shallow) model with two convolutional, one final linear, and two large pooling layers against a six-layer MLP. Deneu et al. [17] follow up and investigate practices found to work well in the DL community: they employ an Inception-v3 [80], one of the main successful CNN architectures for computer vision, but on environmental covariates of variable extent (all of size 64 *×* 64 pixels, but with different resolutions ranging from 30m to 1km). Gillespie et al. [26] employ a TResNet [71], a follow-up on the highly popular ResNet [33] on 1m-resolution RGB-NIR aerial imagery over California. They then combine the TResNet’s features with those from a four-layer MLP receiving 1 km-resolution climatic covariates as input. In both studies, the CNNs (Inception-v3 and TResNet) are pre-trained on the ImageNet classification challenge [19] and then adapted and trained (fine-tuned) on their respective SDM dataset. Such procedure (pretraining on an auxiliary dataset followed by fine-tuning) is known as transfer learning and almost ubiquitous in many DL applications, as it reduces the amount of training data required and makes training DL models on many (relatively small-scale) datasets actually possible.

### 1.3 Assessing the potential of Deep Learning SDMs

Unlike traditional (“shallow”) methods, DL not only requires large training corpora, but can scale along in learning capacity and ability to capture complex relationships as dataset sizes increase. The sub-family of convolutional neural networks (CNNs) inherently takes spatial context into account and can learn to compute the most suitable features (*i.e.*, recombinations based on input covariates) at the right locations around a data point, enriching the information available to inform habitat suitability scores. The modularity of DL further allows for high flexibility in model architectures, which provides three advantages that make them highly suitable for large-scale, multi-species SDM scenarios: first, DL models can be designed to ingest multiple input covariates of arbitrary size and modality simultaneously. This enables DL-based SDMs that can reason not just on bioclimatic point covariates, but also spatial surroundings of different extents and different characteristics, sequential (time series) data, and even semi-structured inputs like textual descriptions of a habitat, all at the same time. Second, DL models can likewise predict multiple targets at the same time and with a common base set of layers. Applied to an SDM context, this means that DL models can be set up as (implicit) joint SDMs, predicting multiple species at once with one common set of base features, possibly borrowing strengths across species [24]. Third, it is comparably straight-forward to train DL models on incomplete observations; species whose presence or absence status is not known for a particular location can simply be ignored by masking during training, a means not commonly possible for conventional models.

All this suggests that that DL should be an attractive contender for SDMs in increasingly large-scale data settings. To date, the exploration of these potential benefits of DL for SDMs has, however, remained scarce. Specifically, a formal, large-scale comparison of DL-based compared to existing SDMs in identical settings is missing, despite rapidly growing use of DL in spatial modelling. Hence, the question of whether and when DL-based SDMs offer competitive or even improved predictions at biogeographic scales and what their biases and limitations might be has remained unanswered. Here, we set out to close this gap and perform a continental assessment of the absolute and relative performance of DL-based SDMs. Specifically, we use DL for predicting the encounter likelihood for 2,299 species of three vertebrate and two invertebrate taxa occurring in North America. We carefully include design choices and modelling heuristics of DL methods and compare architectures specifically designed for the task at hand, including a (more traditional) multi-layer perceptron (MLP) that predicts from point observation data alone, and a CNN that takes spatial neighbourhood into account. We compare them against baselines commonly used in ecology, Maxent [66] and random forest [12], keeping all settings and conditions the same wherever possible, and adjusting and discussing experimental design choices where needed to properly account for peculiarities encountered when using DL models like CNNs. We further provide an evaluation of among-species variation in performance. To our knowledge, this is the first continent-scale, *apples-to-apples* comparison study of DL against established approaches to predicting species distributions.

## 2 Material and Methods

### 2.1 Study Area and Data

All analyses are conducted over the North American continent, within a rectangular box with dimensions (179.99°W, 42.68°W, 10.53°S, and 87.11°N). This includes Canada, the United States, Belize, El Salvador, Guatemala, Honduras, Mexico, and Nicaragua.

#### 2.1.1 Environmental Covariates

We calculate environmental covariates from 17 raster layers at 1 *km*^2^ resolution, all aligned to the same extent, origin, and coordinate reference system (WGS 84). We use various climatic, topographic, vegetation-related, and pedologic covariates; details are provided in Table 1. For the point-based models (Maxent, random forest, MLP), we extract pixel values directly at the centre of each observation, resp. pseudo-absence (see below); in case of the CNN, we extract patches of 64*×*64 pixels around the centre location. We then calculate covariate-wise statistics (average, min, max) for each taxonomic group and all locations (points), resp. a subset of 32,768 randomly sampled locations (patches). Some of the covariates are not defined over aquatic bodies; we replace these pixels in patches with the calculated average. Finally, for all gradient-based learners (*i.e.*, MLP and CNN), we normalise all covariate values in both points and patches by z-score normalisation as follows:

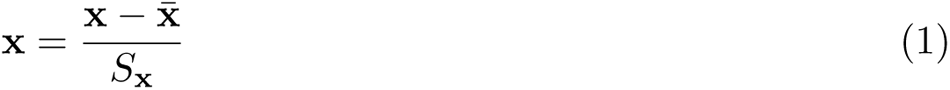

where **x**^-^ and *S***_x_** are the covariate-wise mean and standard deviation as described above, respectively. We do not perform this normalisation for Maxent, which requires higher variation across covariates (we found Maxent not to learn well at all with normalised covariates).

**Table 1:** Covariates used across taxonomic groups. EVI = Enhanced Vegetation Index.

| category | covariate | taxonomic group |  |  |  |  |
| --- | --- | --- | --- | --- | --- | --- |
|  |  | amphibians | ants | butterflies | mammals | reptiles |
| climate | mean annual air temperature [40] | ✓ |  | ✓ | ✓ | ✓ |
|  | temperature seasonality [40] | ✓ |  |  |  | ✓ |
|  | mean temperature of the coldest month [40] |  | ✓ |  |  |  |
|  | annual precipitation [40] | ✓ |  |  |  | ✓ |
|  | precipitation seasonality [40] | ✓ |  | ✓ | ✓ | ✓ |
|  | mean monthly precipitation of the driest quarter [40] |  | ✓ |  |  |  |
|  | accumulated precipitation in growing season days [40] | ✓ |  |  | ✓ | ✓ |
|  | cloud-cover intra-annual variation [86] | ✓ |  | ✓ | ✓ | ✓ |
| topography | topographic ruggedness index [2] | ✓ | ✓ | ✓ | ✓ | ✓ |
|  | topographic wetness index [55] | ✓ |  |  |  | ✓ |
| productivity | mean canopy cover [76] |  | ✓ |  |  |  |
|  | spatial CV of EVI [82] | ✓ |  |  | ✓ | ✓ |
|  | mean summer EVI [21] | ✓ |  | ✓ | ✓ | ✓ |
|  | mean winter EVI [21] | ✓ |  | ✓ | ✓ | ✓ |
| soil | soil silt content [68] | ✓ | ✓ | ✓ | ✓ | ✓ |
|  | proportion clay [68] | ✓ | ✓ | ✓ | ✓ | ✓ |

#### 2.1.2 Species data

Species occurrences for all five taxonomic groups have been retrieved from GBIF [81]; sightings for ants additionally comprise data from AntWeb [65] and Global Ant Bioinformatics Information (GABI; [29]). Occurrence records were subject to various cleaning procedures, including removal of duplicates and spatial/temporal outliers with CoordinateCleaner [94] as well as thinning to limit the number of records per species to one for each square kilometre. We further discarded species with less than six observations in total. A histogram of observations per species and taxonomic group can be seen in Figure S1.

Our models require at least background samples (Maxent), resp. absences (all other models). Since obtaining true absences is difficult for most taxonomic groups and species studied, we sampled pseudo-absences. The heuristic to do so is delicate and can be detrimental to the performance of models; moreover, there is little consensus on the best strategy [5, 67]. In this study, we opt for sampling a constant number of pseudo-absences from within each species’ domain as follows: we first obtain hand-drawn expert range maps as polygons for all species; from Roll et al. [72] for reptiles, from IUCN [35] for amphibians, from Wilson et al. [86] and Map of Life [54] for mammals.

Data for all taxa and species is aggregated and visible on Map of Life^1^ We then calculate the species domain as the union of the expert range map and species-specific observations with a 200 km buffer. Then, for each species, we sample 2,000 pseudo-absences from within the species domain and 0.2 degrees (around 22 km) away from the continent’s coastline (to avoid samples off terrain and with too many undefined covariate values over ocean bodies). Since some species domains are very small, we repeat sampling until we reach 2,000 absences or have tried sampling 24 times; nonetheless, some species thus receive less than the targeted number of absences. The final number of data points per split and taxon group is listed in Table S2.

#### 2.1.3 Data Splitting

We experiment with different ways of data splitting (Figure S11): *random* assigns observations and pseudo-absences uniformly randomly into splits; *spatial* divides the study area into vertical stripes with width according to split fractions; *cluster-spatial* assigns observations by aggregating clusters using k-means on the points’ spatial coordinates. Finally, *ecoregions block-spatial* is the method used for all main experiments: we first bin all data points into square cells of 1*^◦^* lat/lon on a homogeneous grid across the entire study area. Then, we assign each cell to its corresponding North American Ecoregion [61] by testing its four corners (minus 0.001*^◦^* of buffer) and centre location and retaining the mode. For each ecoregion, we iterate through its overlapping grid cells in random order and assign data points of entire cells to either of the three splits so that the number of points match the predefined percentages of 70, 10, 20 for the training, validation, and test set, respectively.

### 2.2 Prediction Models and Training

#### 2.2.1 Multi-Layer Perceptron

The architecture of the MLP is illustrated in Figure 1 top and consists of four primary blocks of a linear (fully-connected) layer, batch normalisation [36] and rectified linear unit (ReLU) activations with negative slope (leak) of 0.01. The first fully-connected layer maps from the *C*-dimensional input point covariates to a latent 512-dimensional vector; this dimensionality is kept for all subsequent blocks (*i.e.*, all but the final fully-connected layers provide a 512-dimensional vector as output). We add dropout [78] with probability of 0.4 before the final fully-connected layer. We experimented with different positions, numbers of dropout layers and dropout rates and empirically found this setup to yield the best accuracy and mitigation of occasional overfitting. A final fully-connected layer maps from the 512-dimensional penultimate latent feature vector to the *N* species, which get turned into pseudo-probabilities with a sigmoid activation.

**Figure 1:**
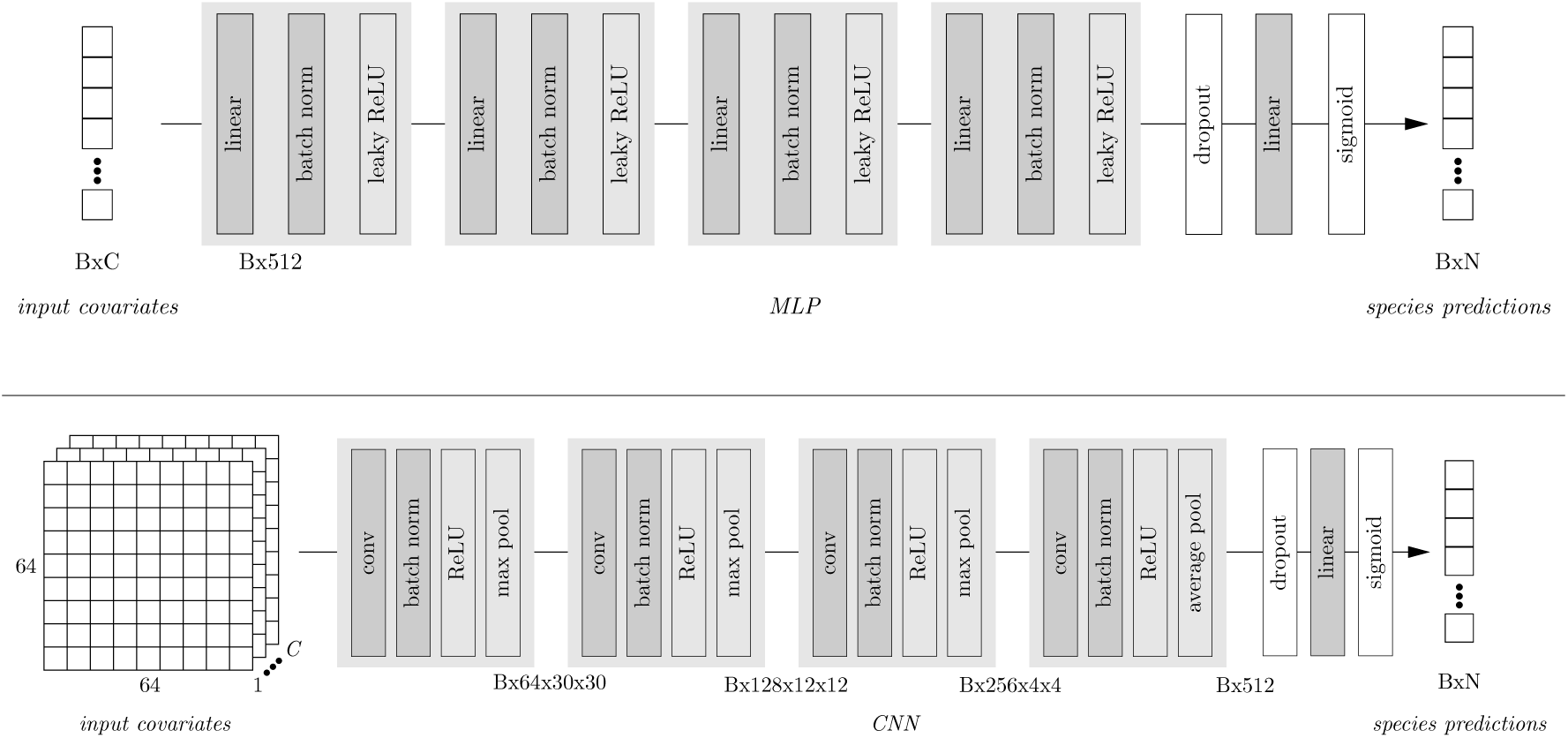
Architectures of the employed MLP (top) and CNN (bottom). *B*: batch size; *C*: number of covariates/channels; *N*: number of species. For the MLP the number of hidden tensor sizes is kept constant at 512 throughout the model; for the CNN intermediate tensor sizes are indicated at the end of each block of layers in format *B × C*[*×W × H*].

As an objective function, we employ the common binary cross-entropy loss:

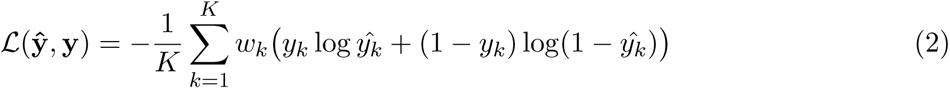

where *y*^*_k_* and *y_k_*are the predicted pseudo-probabilities and ground truth label in *{*0, 1*}* for the *k*th out of *K* classes (species), respectively. *w_k_* is a species-specific scalar weight for the loss term, which we set as follows:

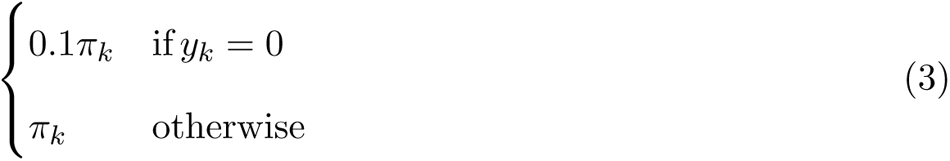

where *π_k_* is the relative frequency of observations to total number of data points per species in the training set:

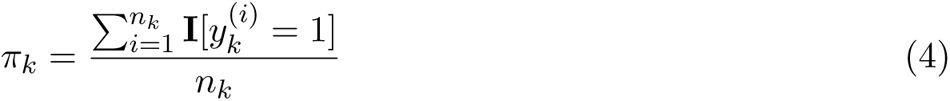

with *n_k_* denoting the total number of training points for species *k* and **I**[*·*] being the indicator function, returning 1 if the condition inside the square brackets is true and 0 otherwise. In other words, we scale the loss of each model prediction element according to the inverse frequency of observations in the training set for each species, and then downscale it again by a factor of ten if it is a pseudo-absence. We tried many different weight regimes, but empirically found these values to work best overall.

Our label set is incomplete in that both occurrences and pseudo-absences are only defined per species, not for the entirety of the model predictions. Hence, we employ binary masks that only calculate the loss for the appropriate model outputs given the correct species. Since the binary cross-entropy loss does not compare across species, it allows the model to predict presence likelihoods independently for multiple species. As in the name of the loss, it minimises the entropy across predictions; *i.e.*, it encourages the model to predict maximal probabilities of one (definitive presence of species) or zero (absence). Although our label space is not fully supervised due to unconfirmed absences, we empirically found this loss to converge well.

We then train each MLP with stochastic gradient descent for 100 epochs (iterations over the full training set) with learning rate of 10*^−^*^3^, reduced by a factor of ten at epoch 50. We also employ weight decay (an *l*1-/lasso-regularisation of the model parameters to prevent overfitting) with strength of 10*^−^*^3^. Momentum, a strategy to channel gradients over subsequent parameter updates, is disabled. We experimented with multiple variations (no learning rate reduction, ten times stronger/weaker weight decay, momentum of 0.9), as well as Adam [43] as a different optimiser, but found the mentioned set of parameters to result in the best convergence of the model. In general, we noted that practices to accelerate learning, such as higher learning rate and momentum, led the model to converge faster, but to less optimal states with lower prediction performance. The batch size is set to 32, which nowadays is very low for such a lightweight setup [32, 39], but again led to the best performance in comparison to values of up to 8,192. Although unprovable in our setting, we hypothesise that smaller batches are more effective as they better draw out subtle variations across covariates and species, with the model better adjusting to individual data points and gradients not cancelling each other out.

#### 2.2.2 Convolutional Neural Network

Similar to the MLP, we design a CNN with four feature extraction blocks (Figure 1 bottom), but gradually scale up the number of hidden features towards 512 at the end. Each block is a sequence of 2D convolution (kernel size 3 *×* 3), 2D batch normalisation, ReLU and 2D max pool, except for the final block, which contains a 2D adaptive average pooling layer instead of a 2D max pool to reduce any spatial tensor to a single, 512-dimensional feature vector. To mitigate effects of spatial autocorrelation, each but the last 2D convolution layer is both strided (2 *×* 2) and dilated (2 × 2). Stride controls the spacing between locations in width and height where the convolutional filter is applied; dilation (also known as *à trous* convolution; [13]) defines the spacing, resp. spread of the locations within the filter itself. Increasing both expands the area considered by the layer and allows it to skip immediately neighbouring locations. Each max pooling layer (size 2 × 2) further provides a mechanism for some translation invariance. We train each CNN with exactly the same hyperparameters as the MLP, as well as additional data augmentation in form of horizontal and/or vertical flips, and up to three 90*^◦^* rotations of the input tensor, each with 50% probability. This forces the model to become invariant to orientation of spatial features and also increases the training set size for better exposure of the model to the underlying data distribution. We also experimented with minor value jittering by adding a bit of normalised noise to the inputs, but this only resulted in model performance degradation.

We implement both MLP and CNN in PyTorch ver. 2.1.1 under Python ver. 3.8.18 and train and run all models on a high-performance cluster node with an NVIDIA Tesla V100 graphics processing unit with 32 GB of video memory.

#### 2.2.3 Random Forest

As a first baseline, we employ random forests [12]. Although random forests can be employed as multi-class predictors, just as DL models, they cannot be used in a setting where classes are not judged against each other. Hence, we fit a separate model for each species and validate and test each model independently. We first perform a hyperparameter search based all mammal species, varying the number of trees as well as the minimum leaf size, both of which allow for control of (over-) fitting. Table S3 lists average AUC_ROC_ and mAP scores on the validation set. We then train random forests with 1,000 trees and minimum leaf size of 11 for the main comparisons.

#### 2.2.4 Maxent

As second baseline, we employ Maxent [66], which fits a distribution across all candidate locations while *(i.)* making sure the distribution matches the averages of the features (covariates) over the observations, and *(ii.)* keeping it as uncertain as possible everywhere else where no a priori information is provided. This latter aspect corresponds to maximising the (Shannon) entropy of the distribution [37].

Maxent is species-agnostic; we therefore fit one model per species. To do so, we use R ver. 4.2.0 and employ the maxnet package (ver. 0.1.4). We do not normalise the input covariates. Maxent offers a range of built-in feature calculation methods on top of raw covariates, including linear (unchanged), hinge and quadratic terms, as well as interactions (product of covariates). A second hyperparameter of the model is the overall regularisation strength (Maxent applies an *l*1 norm as default regularisation scheme with strength modulation depending on the features). Results on the validation set for mammals of different feature sets and regularisation strengths are shown in Table S4. In the end, we use all four features and a constant regularisation factor of 2.0 for all experiments.

### 2.3 Ensemble

To simulate and evaluate a practical approach, we also create an ensemble of all four above-mentioned models as is common in the SDM literature [3]. We do this in a post-hoc selection manner: for each species in each taxon group, we obtain predictions with all four models on the validation set and then use these to calculate model-specific AUC_ROC_ scores. We then select, for each species separately, the one particular model that yields the highest AUC_ROC_ scores on the validation set, and use it to produce the predictions on the final test set. For any species with no samples in the validation set, we use the training set for model selection instead.

### 2.4 Model Validation

We assess performances of trained models on the test set of observations and pseudo-absences for each species and each taxonomic group. In detail, we report the Area under the Receiver Operating Characteristic curve (AUC_ROC_) and mean Average Precision (mAP) evaluation metrics. They are calculated using all unique suitability scores predicted per model. We use scikit-learn ver. 1.3.0 to calculate all performance metrics.

### 2.5 Prediction Post-Processing

Some applications (*e.g.*, Figure 5) require converting continuous suitability score to binary predictions via thresholding. To do so, we employ the commonly used SPS threshold, also known as max TSS [1], which is the arg max of sensitivity and specificity:

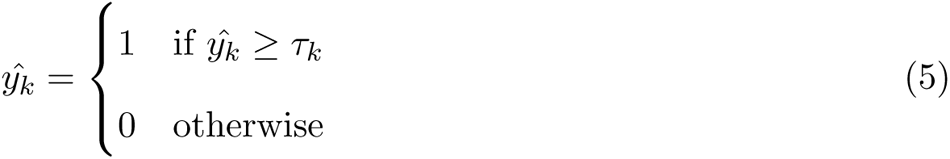

*τ_k_* is the SPS threshold for species with ordinal *k* and calculated as follows:

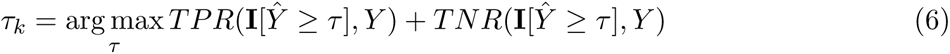

where *τ_k_* is sampled from all distinct confidence thresholds as predicted by the model, and *TPR* and *TNR* are the true positive and true negative rates (sensitivity and specificity), respectively:

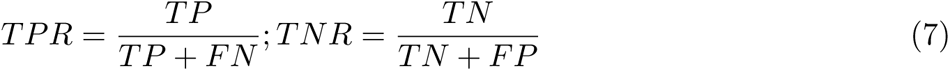

where *TP*, *FP* and *FN* are the number of true positives, false positives and false negatives, respectively. All SPS thresholds are calculated on the full validation set for each model and species separately.

In addition, we produce maps based on a grid of 10 *km*^2^ resolution, with covariate values sampled from the centre of each location (resp. patches around each). All data get normalised the same way (for MLP and CNN) and with the same values as the training data; no data augmentation is applied. For the species richness maps shown in Figure 5, we additionally limit predictions per species to locations lying within its domain, *i.e.*, we set predictions for a particular species to zero for all pixels that lie outside the calculated species domain.

### 2.6 Performance Predictors

Finally, we conduct a formal statistical assessment of putative drivers of the relative performance of DL models at species level. We fit generalised linear models on the delta AUC_ROC_ (Maxent – MLP and Maxent - CNN) as target, and predictors as follows: log number of observations per species (*log no. obs.*), log size of the species’ range map area in *km*^2^ (*log range size*), average latitude of per-species observations (*avg. lat.*). For a subset of the three taxon groups *mammals*, *amphibians*, and *reptiles*, we further include species IUCN threat status as predictor (*threat status*), both ordinal and binarised into *non-threatened* (statuses “Least Concern” (LC) and “Near-Threatened” (NT)) and *threatened* (“Vulnerable” (VU), “Endangered” (EN), “Critically Endangered” (CR), and “Extinct in the Wild” (EW)). We conduct this specific analysis for those species where threat information is available (n=732); all other models are fitted for all species in all five taxon groups (n=2,167). All regression models are fit and metrics reported in Python with statsmodels ver. 0.14.1.

## 3 Results

### 3.1 Overall Modelling Performance

Across all 2,299 species five taxa analysed in comparable fashion and with taxon-specific DL models, we find a broadly similar performance between the two DL approaches (MLP, CNN) and the two traditional, shallow models (Maxent and random forest) (Table 2). Area under the Receiver Operating Characteristic curve (AUC_ROC_) mean (standard deviation in brackets) across all five taxa combined amounts to 0.78 (0.14) for the taxon group-specific MLP, −0.03 (+0.03) compared to Maxent, and 0.76 (0.14) for the CNN, −0.05 (0.00) compared to Maxent. Histograms are less right-skewed and have a lower kurtosis for MLP and CNN, compared to the shallow models (Figure S2), and CNNs stand out for causing a particularly large number of species with low AUC_ROC_.

**Table 2:**
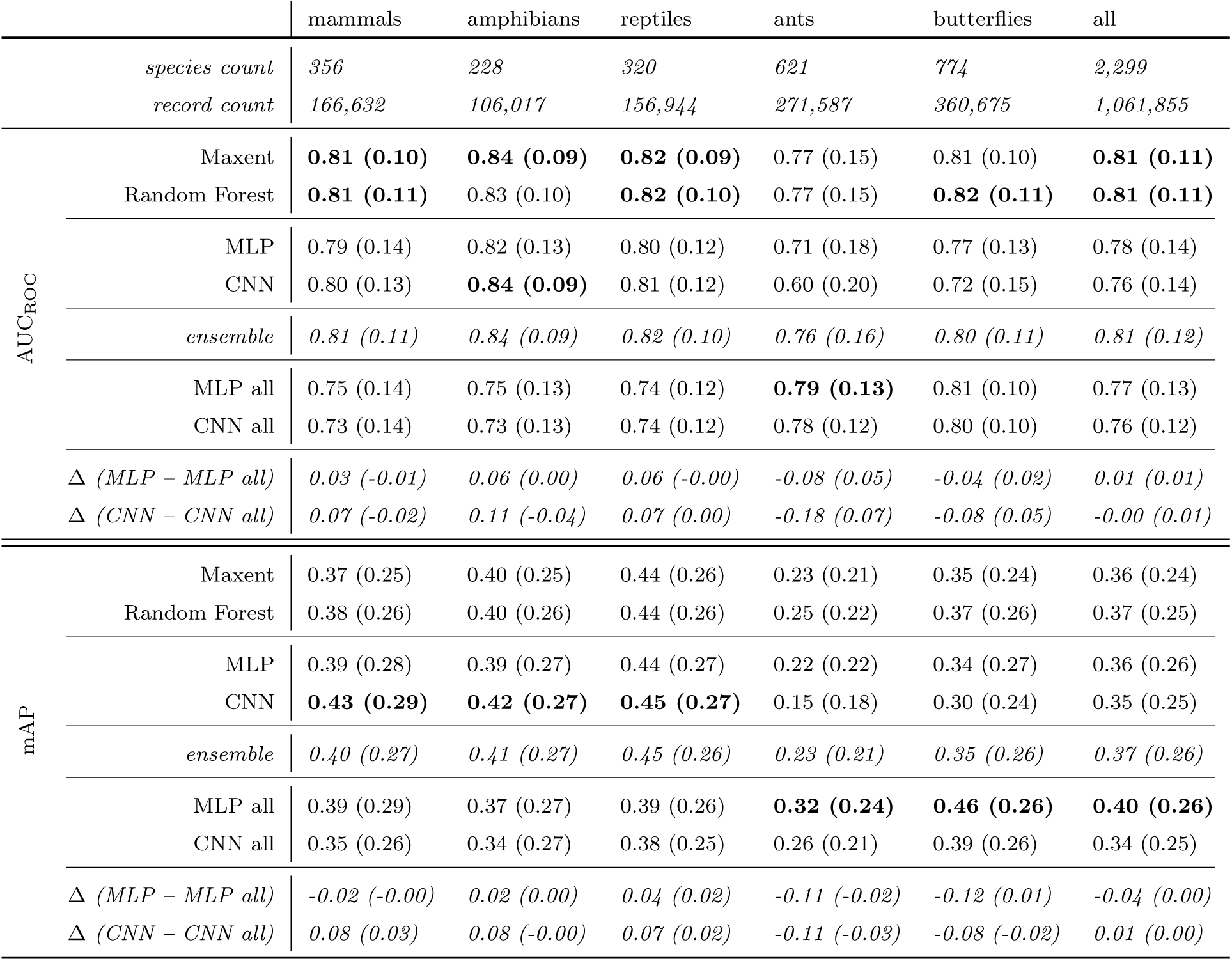
Area under the Receiver Operating Characteristic Curve (AUC_ROC_) and mean Average Precision (mAP) scores on the held-out test set given as average (standard deviation) across all species for each taxon group and model. The two Deep Learning (DL) models “MLP” and “CNN” were trained for each taxon group separately, whereas “MLP all” and “CNN all” were trained on the full set of 2,299 species. “ensemble” is a mixture where one of Maxent, Random Forest, MLP, and CNN is chosen, based on highest AUC_ROC_ score on the *validation* set (hence scores on the test set may be lower). Bold numbers report the per-taxon group highest performance. All figures, including record counts, are reported for the held-out test set.

We find that for the DL models in particular performance is highly affected by choice of data splitting procedure. Using a typically employed *cluster-spatial* approach (observations are assigned to train/validation/test splits as entire spatial clusters, pseudo-absences are distributed randomly; see Materials and Methods section) results in a highly inflated AUC_ROC_ scores, particularly for the CNN, scoring an average AUC_ROC_ of 0.87 against 0.77 of Maxent (Figure 2). This can be attributed to the CNN’s explicit utilisation of spatial neighbourhood, which causes excessive information leakage across splits in the “cluster-spatial” approach (see below). Unless noted otherwise, all results pertain to models trained and evaluated under the more conservative *ecoregions-block-spatial* split.

**Figure 2:**
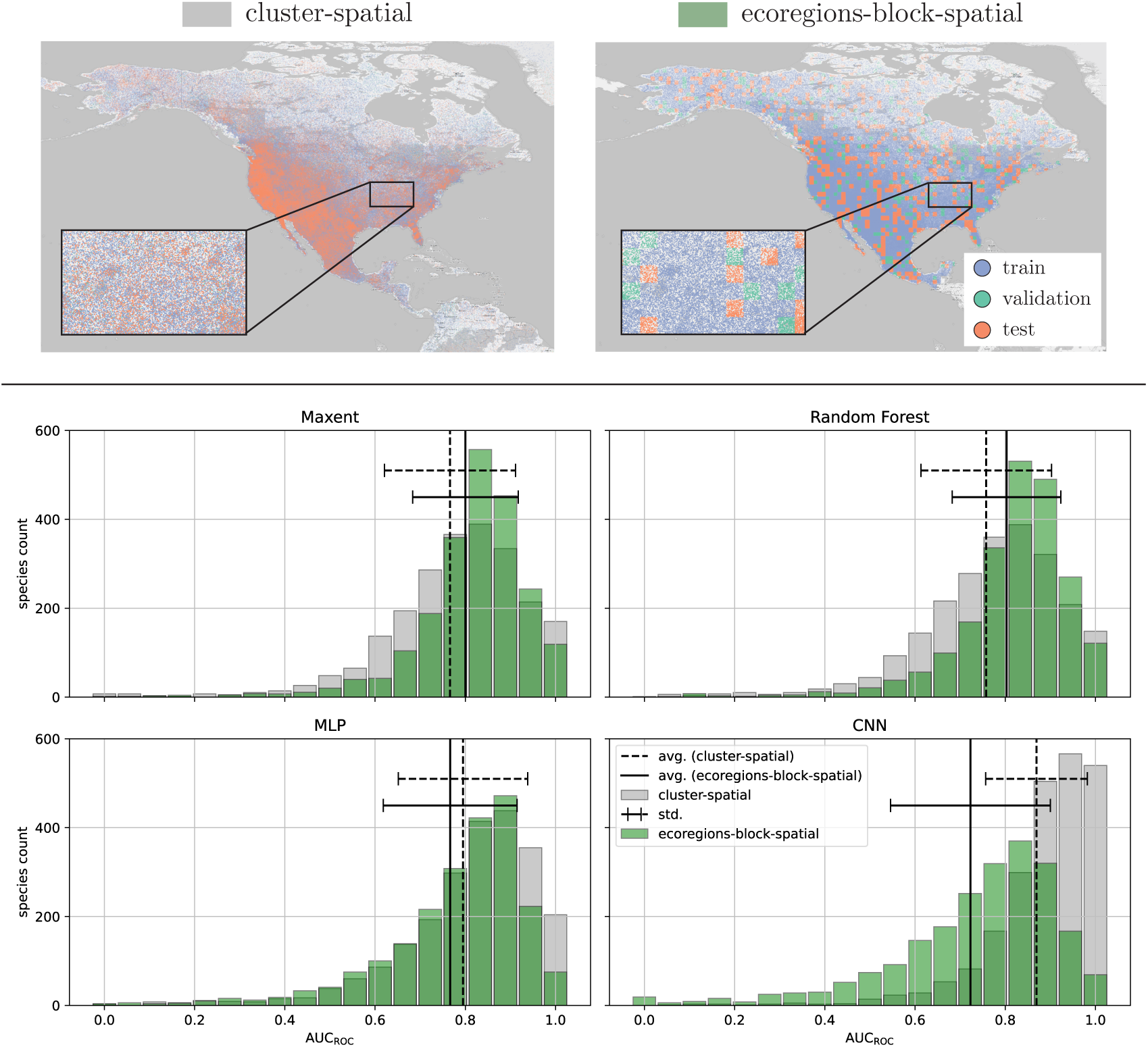
Performances under different data splitting procedures: cluster-spatial (grey) and ecoregions-block-spatial (green). The top row shows data point assignments in geographic space; the four panels below display histograms of AUC_ROC_ on the test set for all four models, combined over all five taxonomic groups (Maxent, Random Forest: one model per species; MLP, CNN: one model per taxon group). Also shown are average and standard deviation AUC_ROC_ values for cluster-spatial (dashed) and ecoregions-block-spatial (solid).

Among the five taxa, taxon group-specific DL models (“MLP”, “CNN”) perform similarly or inferior compared to the top-performing shallow models in terms of AUC_ROC_ (−0.17 to 0.00 difference), and similarly to slightly better in mean Average Precision (mAP) (−0.10 to 0.05 difference).

DL models are relatively weaker in ants (−0.06 AUC_ROC_/+0.03 mAP) and butterflies (−0.05/-0.03), the most species-rich taxa, and mostly similar or even slightly superior in reptiles (−0.01/0.01), amphibians (0.00/0.02), and mammals (−0.01/0.05). Figure S2 shows additional nuances and highlights a strong heterogeneity of model performance at species level.

A second series of DL models trained over all species and all taxa (*i.e.*, one model run instead of five, “MLP all” and “CNN all”, Table 2) generally shows inferior performance. Specifically, the differences to existing SDMs seem opposite to those seen for taxon-specific DL-SDMs—both DL models trained on all data fall behind in all cases where their taxon group-specific counterparts excel (amphibians, mammals, reptiles), but pull ahead for ants and butterflies where taxon-specific models perform worst.

### 3.2 Relative DL-based SDM performance across species

The performance of DL-based and existing SDMs varies strongly among species (Figure 3 A). While for most species both MLP and Maxent perform well, many cases exist where strong performance in one coincides with poor performance in the other. We next assess the putative drivers of these relative performance differences and specifically evaluate the correlates of high MLP compared to Maxent performance (Figure 3, Tables S5 and S6). We find that relative performance varies strongly by geographic range size: MLP is inferior to Maxent for narrow-ranged species (Figure 3 B). A similar trend is found for sample size, with MLP performance exceeding Maxent only for species when many observations are available for model training (Figure 3 C). Both effects remain significant even as one is statistically accounted for, despite the variables’ collinearity (*R*^2^ = 0.94, *p* = 0.00). Average latitude of observations has a lesser, but still significant effect (Tables S5 and S6), with slight bias towards the DL models performing better for species in more northern regions. The strongest model includes all investigated predictors (latitude, log no. observations, log range size) together for both MLP and CNN (lowest Akaike Information Criterion values). Finally, correlations between IUCN threat status and performance for a subset of species and taxon groups where status is known are significantly negative in case of the MLP, indicating inferior performance for threatened species compared to Maxent (Table S5). Effects were non-significant for the CNN (Table S6).

**Figure 3:**
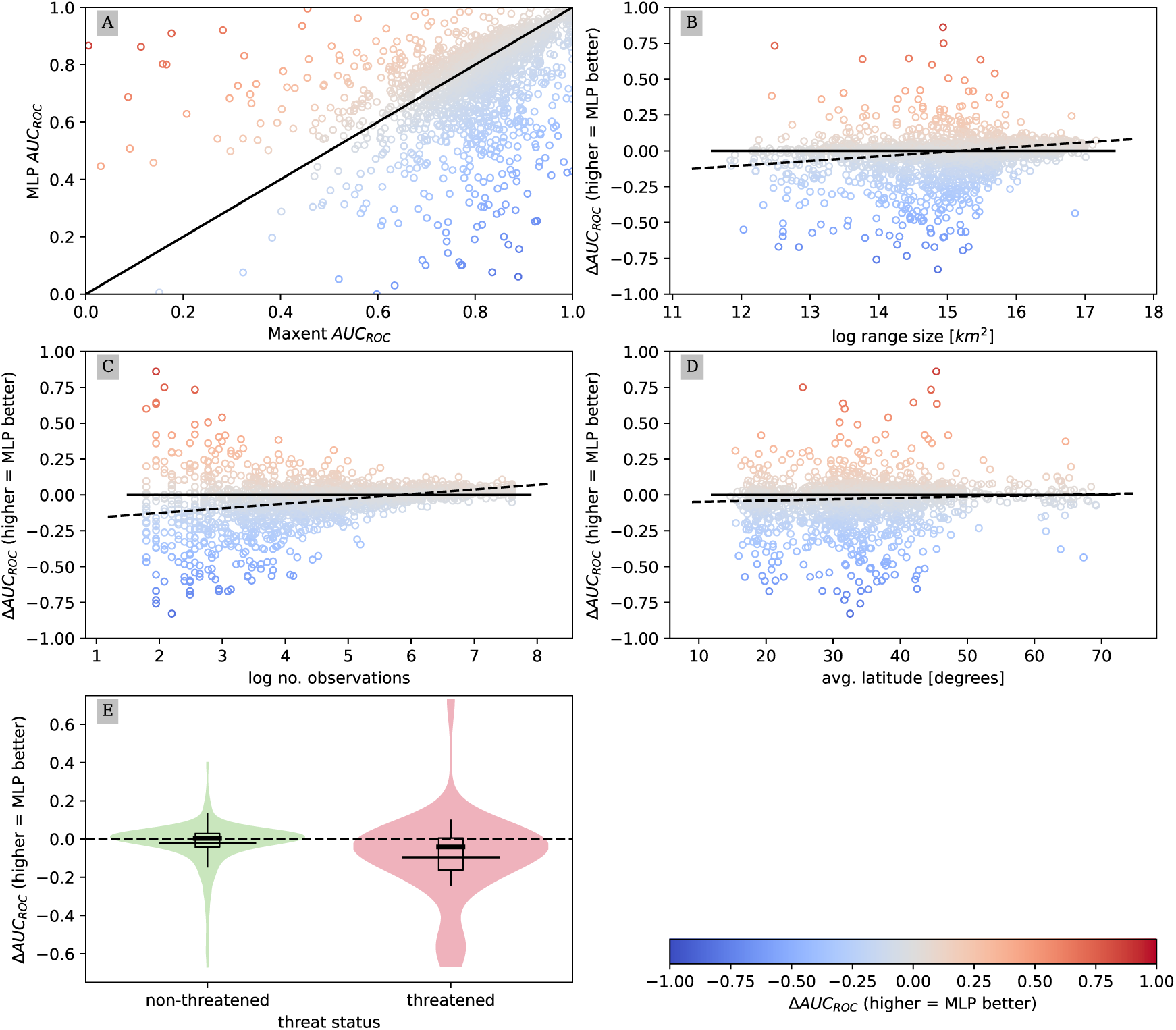
Plots of prediction performances and putative drivers across all five taxon groups: MLP against Maxent AUC_ROC_ (A), delta AUC_ROC_ (MLP – Maxent) against log range size (B), log number of observations (C), average latitude of observations (D), and threat status (E), where “threatened” includes species with IUCN threat status “Vulnerable” (VU) or higher. Also shown are 1:1 (solid) and GLM regression lines (dashed).

DL models are generally known for requiring large amounts of training data; an effect of sample size on model performance is therefore expected. Our results nonetheless uncover important nuances among model types. Notably, a focused analysis into the issue for mammals highlights the superior performance of Maxent up to ca. 40 data points that is followed by relatively slightly smaller

AUC_ROC_ values above a sample size of ca. 150 (Figure 4). Accounting for number of training points per species, both DL models consistently outperform the shallow baselines in AUC_ROC_, mAP, true skill statistic (TSS) and Cohen’s kappa score for all species with around 200 training points or more (Figure 4 and Figures S5-S8).

**Figure 4:**
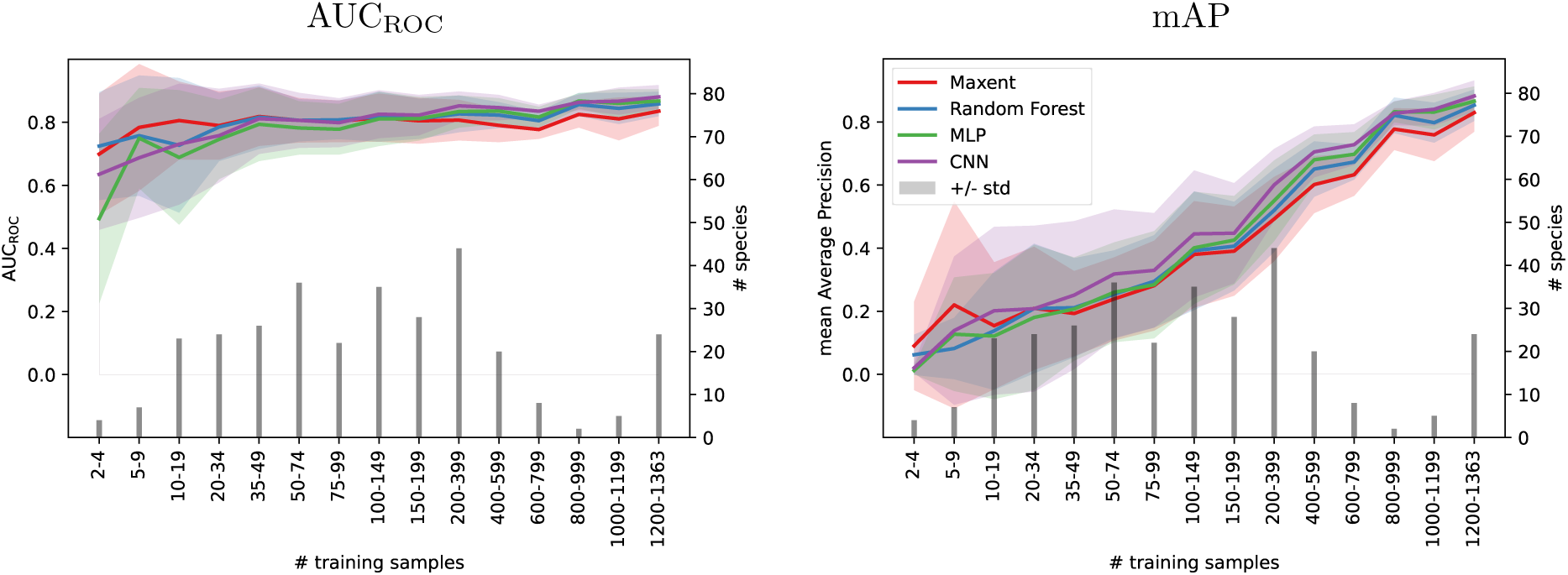
Mean AUC_ROC_ (left) and AP (right) for *mammals* and all models. All values are plotted with their standard deviation (shaded area; left y axis) against species binned by number of training samples (bars at bottom; right y axis).

### 3.3 Comparison in geographic space

A comparison of species richness maps predicted for mammals (Figure 5) reveals that both DL models predict suitability scores that are less spread out and more spatially concentrated than Maxent, exhibiting stronger and narrower peaks. All models closely match a peak of mammal species richness in southern New Mexico/southeastern Arizona, with both MLP and CNN showing higher richness in central Mexico than Maxent, which is only partially reflected in richness estimates based on expert range maps.

**Figure 5:**
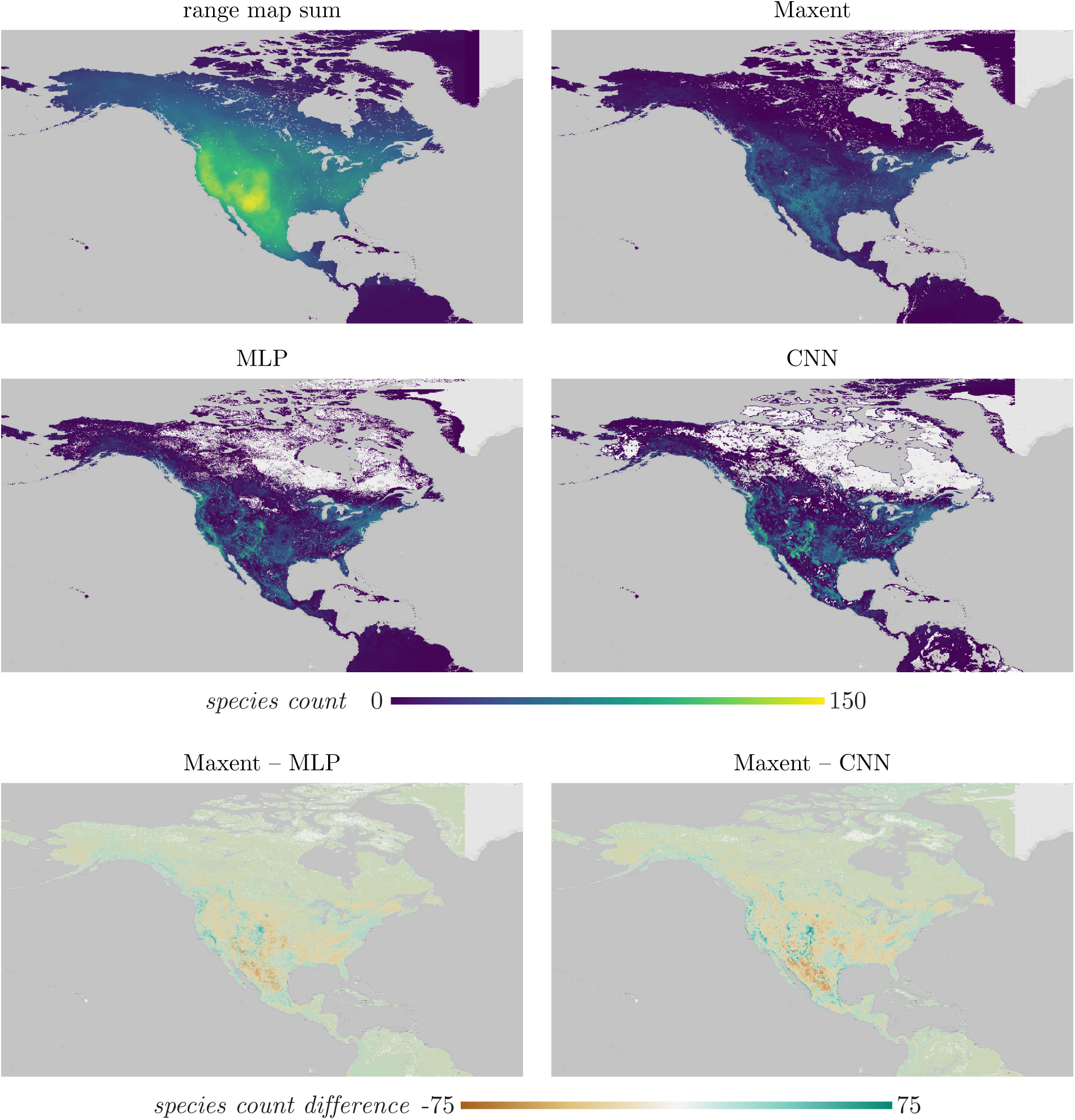
Comparison of richness patterns based on the aggregation of range maps and by thresholded model predictions of the two DL models (MLP, CNN) and Maxent. The bottom row shows the absolute difference between Maxent and the respective DL predictions. For this display, model predictions are limited to the respective range map boundaries for each species (predictions outside are set to zero prior to summation).

**Figure 6:**
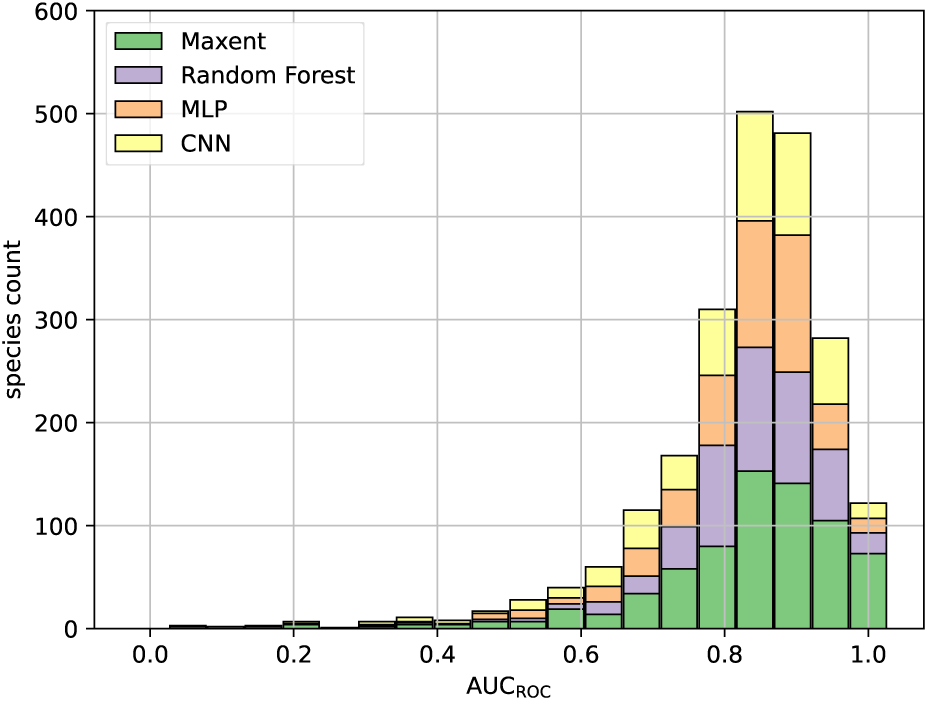
AUC_ROC_ histogram for the ensemble across all five taxonomic groups, with bins stacked regarding selected model per species.

### 3.4 Combining existing and DL-based SDMs via ensemble

Finally, we assess the performance of an ensemble across Maxent, random forest, as well as the taxon group-specific DL models (MLP and CNN), where for each species the model with highest AUC_ROC_ score on the validation set is chosen. For both AUC_ROC_ and mAP, the ensemble either matches individual, best-performing models (mammals, amphibians, reptiles in AUC_ROC_; reptiles in mAP), or trails slightly behind; averaged across all taxon groups, the ensemble is on par with the top-performing Maxent in AUC_ROC_ and behind the combined MLP (“MLP all”) in mAP by 0.03 points (Table 2). An AUC_ROC_ histogram of the ensemble can be seen in Figure S3 top. All four models were selected across all five taxonomic groups, with Maxent selected for 711, random forest for 499, MLP for 485 and the CNN for 472 species (6). Highest absolute contributions occurred for species with test set AUC_ROC_ scores of around 0.84 (Maxent, random forest, CNN), and 0.89 (MLP). Scatter plots of per-species test set AUC_ROC_ against number of observations of the respectively selected model within the ensemble (Figure S4) reveals a partitioning effect of selected models, with Maxent performing best for species with low observation count, random forest being selected for slightly more observation-rich species, and both MLP and CNN exhibiting high spread yet being selected for most of the species with maximum number of observations (2,000), apart from butterflies (where random forest outperformed both MLP and CNN in some cases).

## 4 Discussion

### 4.1 Dataset Size and Model Generalisability

At first glance, the performance of both DL models is directly related to the support of the species, with both models struggling most for species with few observations, but outperforming all other models for data-rich species. This directly corresponds to expectations from the ML literature. The CNN in particular shows the strongest drops in performance, possibly because it contains roughly twice the number of learnable parameters (1.7 to 2 million) compared to the MLP (900,000 to 1.2 million). Figure 4 indicates that around 200 observations or more per species are required for either DL model to outperform their shallow counterparts in AUC_ROC_; similar numbers apply to the other taxonomic groups. One ecological framework to improve an SDM’s modelling ability of data-deficient species relies on the concept of “borrowing strength”, where species’ niche parameters are partially inferred from other species that are related in some way [24]. Various approaches have been proposed to define this relatedness, such as phylogenetic distances [57] or trait similarities [56], both of which require auxiliary data. In turn, DL methods as employed here model species jointly by design by sharing all but the final layer’s weights across species. While we do not include auxiliary information that establishes explicit relationships across species, we can observe some degree of joint information exploitation in the DL models: both MLP and CNN increase in performance and sometimes even outperform the shallow models for taxa with many species but few observations once information from other taxa is included; see the increase in performance for ants and butterflies of “MLP all” and “CNN all” over their single-taxon versions. For single-taxon models, the regression analysis (Table S5) shows that taxa with high species count correlate the most (negatively) with MLP AUC_ROC_. The taxon group ants, for example, thus seems to be insufficient in size and/or covariate diversity to benefit DL models. With a machine learning lens, this possibly translates to better generalisation ability of these models thanks to exposure to more diverse parts of the underlying species and covariate distribution, once a critical mass of training points has been reached.

The performance gap between data-rich and -poor species might be reduced by drawing on various directives from the machine learning literature. A first directive is the *long-tailed distribution problem* (few species with many observations and many species with few observations), which has been subject to extensive research in the machine learning community [50]. DL models generally focus on the most abundant classes (species) with priority, as this results in the best strategy to reduce the overall training loss; this comes at the cost of reduced performance for the rare classes (the long tail). Various solutions have been proposed to counteract this phenomenon, such as loss re-weighting (which we also apply to some extent here, see Materials and Methods section), augmenting the rare classes with synthetic examples [7], memory banks [50], and custom loss functions [91].

The second research directive is *few-shot learning*, which attempts to train DL models on data with many classes having few training examples each. This directly corresponds to our setting, in particular ants and butterflies, which include many species with few observations. Example methods are prototypical networks that learn representative instances (prototypes) for each class [77], as well as meta-learning, which attempts to train a DL model to be able to adapt quickly across combinations of classes at hand [25]. Our DL models generally work well for species with around 200 training points or more. Given that “low observation count” oftentimes refers to single-digit numbers in ecology, this still implies a huge gap in DL model applicability. Hence, research on such heuristics that address high imbalances and small dataset size regimes is paramount to render DL palatable for typical ecological settings.

### 4.2 Data and validation biases

As DL is no silver bullet, its performance gets influenced by the same biases emerging from the dataset as any other model. Besides sample and range map size, this includes latitude (sampling effort diminishes quickly in sparsely populated Northern Canada) as shown in the regression analysis (Table S5 and Figure 3). The SDM literature shows several means of resolving this bias, such as target-group background sampling [67, 11] and occupancy modelling with inclusion of encounter likelihood [4, 20]. Some of these ideas are readily applicable for DL models and first results have been shown [75, 88], but further research is needed to understand effects and provide recommendations in the DL SDM space.

Another, rather problematic bias on prediction model evaluation in general is information leakage across folds or splits, *i.e.*, signals from the test set data points exposed to the model during training. Spatial analyses quickly compromise split integrity due to spatial autocorrelation; resolution only lies in clear spatial separation of the splits. This is particularly difficult for SDMs, as all splits must not only be free from leakage, but also cover all expected environmental configurations like ecoregions employed in this study (see Materials and Methods section), and contain enough examples for all species in case of multi-species SDMs like ours. Spatial models like our CNNs unfortunately add yet another level of complexity: they ingest a patch of covariates around the spatial location of each data point and require an even greater minimum distance between locations of different splits. With a base 1 km^2^ resolution and CNN designed for an input patch size of 64 × 64 as in our case, this would require dropping too many training points, which results in strongly compromised model performance and evaluation.

Our analyses of model performance under different splitting scenarios showed highly inflated AUC_ROC_ when a conventional yet compromised split (“cluster-spatial”) was employed, with the CNN in particular scoring up to +17 points compared to the second-best performing model (Figure 2). Results on the finally employed heuristic, “ecoregions-block-spatial”, reveals that this result is strongly biased, with the CNN not performing better. The MLP also shows moderate signs of biases, which most likely is attributable to information leakage across species per location instead of spatial autocorrelation alone. Although ecoregions-block-spatial reduces leakage, it compromises representativeness of species across splits, in particular for species with narrow home range and few observations (Figure S11).

### 4.3 Implications of DL as SDMs in Practice

A spatial distribution map is arguably most useful for species that are under particular scrutiny, such as data-deficient or endangered ones [38]. This unfortunately goes against the performance band of DL models, which dictates that data-rich species (in our empirical study, species with 200 observation points or more) get predicted most accurately. DL methods further pose high computational load (training time took between five hours for a single MLP and two weeks for “CNN all”) and increased implementation complexity.

However, just as DL is not a single model, employing it for a particular application is not a one-stop shop. While we have made careful design considerations and have tuned model hyperparameters by-the-(machine learning) book, we nonetheless have had to make many assumptions and compromises. Below is a subset of considerations that merit further investigation in this regard:

- Like random forests, our DL models are discriminative classifiers. They therefore assume a fully-observed dataset, which is not the case in our presence-only setting. We still obtain decent results with semi-randomly sampled pseudo-absences and carefully tuned loss weights, but note that this remains a compromise. [15] proposes and examines multiple ways of training DL models in such a setting, all of which reduce to alterations in loss functions. The more general realm in machine learning that is perhaps most similar to opportunistically sampled species location data is *positive-unlabelled learning* [8], which, to the best of our knowledge, has not yet been applied to DL SDMs.
- Since pseudo-absences have a different meaning and magnitude for Maxent *vs.* random forest *vs.* DL, their sampling strategy and number should be subject to further investigation. Maxent is generally known to require many background samples to be able to properly estimate covariate distributions [5]; in the case of DL, however, too many absences could easily drown the learning signal from the observations [91, 50].
- In theory, there is no requirement for a DL model to be a classifier; instead, one could very well imagine a model designed without a discriminative objective. One such example is the family of deep metric learning [52], which maps data points to a latent feature space and pulls similar points (*e.g.*, of the same species) together and pushes dissimilar ones apart. Theoretically, this requires no absence points at all. We initially tried a model with a deep metric learning objective, but found the high complexity of sampling representative combinations of data points required to calculate the loss prohibitive.

These considerations may lead to increased performance of DL-based SDMs once further investigated. Only then could DL play its full potential for the task of species distribution modelling. As addressed in the Introduction, DL entails a number of promising capabilities that have not even been explored in this basic, apples-to-apples comparison study, including its capability to ingest arbitrary numbers and modalities of data inputs and to provide equally diverse predictions. Some early propositions in this regard have recently been made, such as the inclusion of large language models [74]. Furthermore, it is worth noting that the DL models employed in this study are exceptionally small, compared to present-day architectures employed in other disciplines. We attempted to use a somewhat larger, more common CNN architecture (ResNet-18 [33]) but obtained inferior performance, indicating that the complexity of spatial patterns in environmental covariates at 1 *km*^2^ resolution does not merit larger models. Larger CNNs have been shown to be beneficial for very high-resolution (*i.e.*, 1 *m* and less), imaging covariates [26]. However, using such covariates is not (yet) common practice for SDMs and the interpretation of their type and magnitude of contribution to predicting habitat suitability currently remains questionable.

From a machine learning point of view, improvements in modelling capabilities will likely lead to higher predictive performance, probably surpassing existing SDMs at some point and overthrowing them once dataset sizes become too prohibitive for shallow models to handle. However, we argue that SDMs should bring more to the table than high predictive power: being able to make *inference* (*i.e.*, understand model processes and potentially uncover new relationships, for example across species), ought to be more important than numerical performance under oftentimes biased data setups. This also reaches beyond the generation of single-species (or stacked) distribution maps for which SDMs are often used, and could particularly benefit analyses at an aggregated level, for example pertaining to dynamics of species assemblages in space and time, or else compositions of communities. In sum, we postulate the full power of DL for species distribution modelling to still be pending, only achievable if machine learning and ecology research disciplines do joint efforts and attempt to understand, and include, each other’s requirements and knowledge in the design of DL-based SDMs.

## 5 Conclusion

As the field of ecology is experiencing a shift towards big data, its modelling approaches need to be able to scale along. DL provides unique opportunities to this end, being not just able to better cope with large datasets, but offering novel ways of understanding species distribution and habitat suitability through its ability to accept complex data modalities, as well as perform arbitrarily nonlinear modelling. Yet, successful implementations of DL as SDMs are still rare and tend to lack extensive validation and investigation of biases and limitations.

We conduct large-scale experiments and present one of the first studies that compares DL with conventional SDM methods at large (continental) scale and in an apples-to-apples fashion. Our results show that DL does not yield the same performance margin as in other areas of application.

DL analyses suffer critically due to dataset biases, leading to exaggerated performance figures for DL-based SDMs, especially spatial models like convolutional neural networks. Beyond these biases, DL employed in the same fashion as conventional SDMs currently does not yet appear to offer superior predictive performance. However, we argue that the full potential of DL for modelling species distributions has not yet been exploited and that the era of DL-based SDMs is still to come. Given targeted research, we project DL SDMs to soon be able to fully play their cards, providing insights that go beyond mere numerical improvements. Being able to advance such directions requires profound expertise in both fields of ecology and machine learning, and thus an in-depth collaboration between disciplines.

## Footnotes

1 https://mol.org/patterns/high-res

